# Miniature Structured Illumination Microscope for *in vivo* 3D Imaging of Brain Structures with Optical Sectioning

**DOI:** 10.1101/2021.12.02.470995

**Authors:** Omkar D. Supekar, Andrew Sias, Sean R. Hansen, Gabriel Martinez, Graham C. Peet, Xiaoyu Peng, Victor M. Bright, Ethan G. Hughes, Diego Restrepo, Douglas P. Shepherd, Cristin G. Welle, Juliet T. Gopinath, Emily A. Gibson

**Affiliations:** Department of Electrical, Energy and Computer Engineering, University of Colorado Boulder, CO, 80309, USA; Paul M. Rady Department of Mechanical Engineering, University of Colorado Boulder, CO, 80309, USA; Department of Physics, University of Colorado Boulder, CO, 80309, USA; Department of Bioengineering, University of Colorado Anschutz Medical Campus, CO, 80045, USA; Department of Neurosurgery, University of Colorado Anschutz Medical Campus, CO, 80045, USA; Department of Physiology and Biophysics, University of Colorado Anschutz Medical Campus, CO, 80045, USA; Department of Cell and Developmental Biology, University of Colorado Anschutz Medical Campus, CO, 80045, USA; Department of Physics, Arizona State University, Tempe, Arizona, 85287, USA

## Abstract

We present a high-resolution miniature, light-weight fluorescence microscope with electrowetting lens and onboard CMOS for high resolution volumetric imaging and structured illumination for rejection of out-of-focus and scattered light. The miniature microscope (SIMscope3D) delivers structured light using a coherent fiber bundle to obtain optical sectioning with an axial resolution of 18 μm. Volumetric imaging of eGFP labeled cells in fixed mouse brain tissue at depths up to 220 μm is demonstrated. The functionality of SIMscope3D to provide background free 3D imaging is shown by recording time series of microglia dynamics in awake mice at depths up to 120 μm in the brain.

## 1. Introduction

To further our understanding of neural circuits and their function, there is a need for new tools that can perform high resolution imaging of neural dynamics in animals performing complex behaviors. This goal has motivated development of miniature lightweight microscopes that can be head-attached, providing unrestricted motion for studies of behaviors such as navigation or socialization. A variety of miniaturized microscopes have been developed for recording neural activity from large populations of neurons involved in a common neural computation. For example, fluorescence microscopy in awake animals expressing genetically encoded Ca^2+^ fluorescent indicators, can provide real-time functional information with single cell specificity within local neural circuits. The widely used UCLA miniscope includes an LED source and miniature camera and is designed with enough spatial and temporal resolution to capture GCaMP fluorescent transients from single cells. However, these systems have a large depth of field and do not provide any 3D structural information. Imaging in 3D imposes further requirements on the microscope that include low aberration optical designs, along with the ability to perform optical sectioning and axial scanning to distinguish structures at different depths in the tissue. This opens new applications in resolving functions of neural circuits in different brain regions, as well as studies of structural cellular changes.

Fiber-coupled miniature microscopes using multiphoton excitation have been demonstrated for recording neuronal activity in brain [1–5] along with the capability of volumetric imaging [2,3]. Multiphoton excitation processes used for imaging in these miniature microscopes have inherent optical sectioning and provide high spatial resolution. However, laser scanning technique requires active mechanical scanning elements that limit acquisition frame rate, in addition to bulky and expensive ultrafast pulsed laser sources. These limitations are partly addressed by single photon widefield miniature microscopes, i.e. UCLA miniscope [6–8], NINscope [9], FinchScope [10], CHEndoscope [11], Miniscope [12] and Inscopix [13], with some versions even providing wireless capabilities [7], multi-site recording [9] and axial scanning [14]. Recent work used a widefield miniscope modified by placing a phase plate [15] or microlens array [16] in the optical detection path which provides additional information for computational reconstruction in three dimensions. However, challenges associated with removal of scattered light, particularly to identify structural features, still exist with these modified 3D miniscopes. There is a need for higher resolution imaging to clearly identify individual neuronal cells in dense tissue volumes, and to image processes of both neurons and non-neuronal glial cell populations in the brain.

The idea of obtaining optical sectioning in conventional widefield microscopes by projecting a single spatial frequency grid pattern with three relative spatial phase shifts to obtain optically sectioned images was experimentally demonstrated by Neil *et al*. [17]. The reconstruction method, based on the square law detection scheme, rejects the zero spatial frequency components which are not attenuated out of focus, while capturing the components corresponding to the frequency of the grid pattern. The fluorescence generated from the grid pattern is imaged most sharply from the focal plane, hence providing inherent optical sectioning using this structured illumination microscopy method. Reconstruction techniques using structured illumination have also demonstrated sub-diffraction limited imaging in biological tissue [18–22], in addition to providing optical sectioning. However, these imaging systems can require aberration corrected high numerical aperture (NA) objectives, which poses a challenge for miniature microscopes where weight and dimensions are critical parameters.

Here, we demonstrate the first fiber-coupled miniature microscope with optical sectioning structured illumination to remove out-of-focus fluorescence and scattered light, that can obtain full 3D imaging with improved contrast in scattering tissue. The miniature microscope includes an active axial scanning element providing volumetric imaging. The structured illumination miniature microscope with 3D imaging (SIMscope3D) uses a digital micromirror device [20] to create the structured illumination pattern that is then relayed to the imaging plane using a coherent fiber bundle [23,24]. The onboard CMOS camera with a 2.2 μm pixel size enables high lateral resolution images free of artifacts from the fiber bundle. The electrowetting axial scanning element provides depth scanning of up to 550 μm into the sample. Using the SIMscope3D we demonstrate optically sectioned high-resolution images up to 220 μm deep in fixed brain tissue labeled with PLP-eGFP, and time series multiplane images up to 120 μm of microglia processes motion in an awake mouse.

## 2. Methods and materials

### Imaging System

A light emitting diode (LED) at 470 nm (Thorlabs EP470S04) was used as the illumination source for our imaging system. The fiber-coupled LED output was collimated using an objective lens (Olympus UPlanSapo 10x/0.4 NA) and passed through an excitation filter (Chroma ET470/24m). The collimated and filtered output from the LED was then incident on the digital micromirror device (Texas Instruments DLP Lightcrafter 6500). The spatial pattern generated on the digital micromirror device (DMD) was focused onto the back focal plane of a microscope objective (Olympus UPlanSapo 10x/0.4 NA) using a 200 mm achromatic lens (Thorlabs AC508-200-A-ML). The patterned beam was then coupled into a coherent fiber bundle (Fujikura FIGH-10-500N, 10,000 cores, 2.9 μm core diameter, and 4.5 μm core-to-core spacing [25]) at the focal plane of the microscope objective.

The pattern generated on the DMD is relayed from the distal to proximal end of the fiber bundle and coupled into the miniature microscope. The SIMscope3D is designed with an NA of 0.3 and a magnification of 2.2X, providing a circular FOV with a 207 μm diameter from the 460 μm active imaging diameter of the fiber bundle. Commercially available achromatic doublets are used in the optical design along with a custom dichroic cube beam splitter (Shanghai Optics) to separate the excitation and fluorescence emission paths. The SIMscope3D utilizes an electrowetting axial scanning element (Corning Varioptic A-25H) which provides up to 550 μm of active axial scanning below the cover slip (Figure S1 and S2). The fluorescence generated at the imaging plane is imaged onto a CMOS sensor (Ximea MU9PM-MBRD) through an emission filter (Chroma ET525/50m) integrated with the microscope. The CMOS sensor has a pixel size of 2.2 μm resulting in a lateral resolution of 1μm/pixel. The total weight of the SIMscope3D was 6.7 g with a measured height of 30 mm. A schematic of the miniature imaging system along with capabilities of optical sectioning and volumetric imaging are shown in Figure 1 (detailed description of SIMscope3D design and assembly is given in Figure S3).

**Figure 1:**
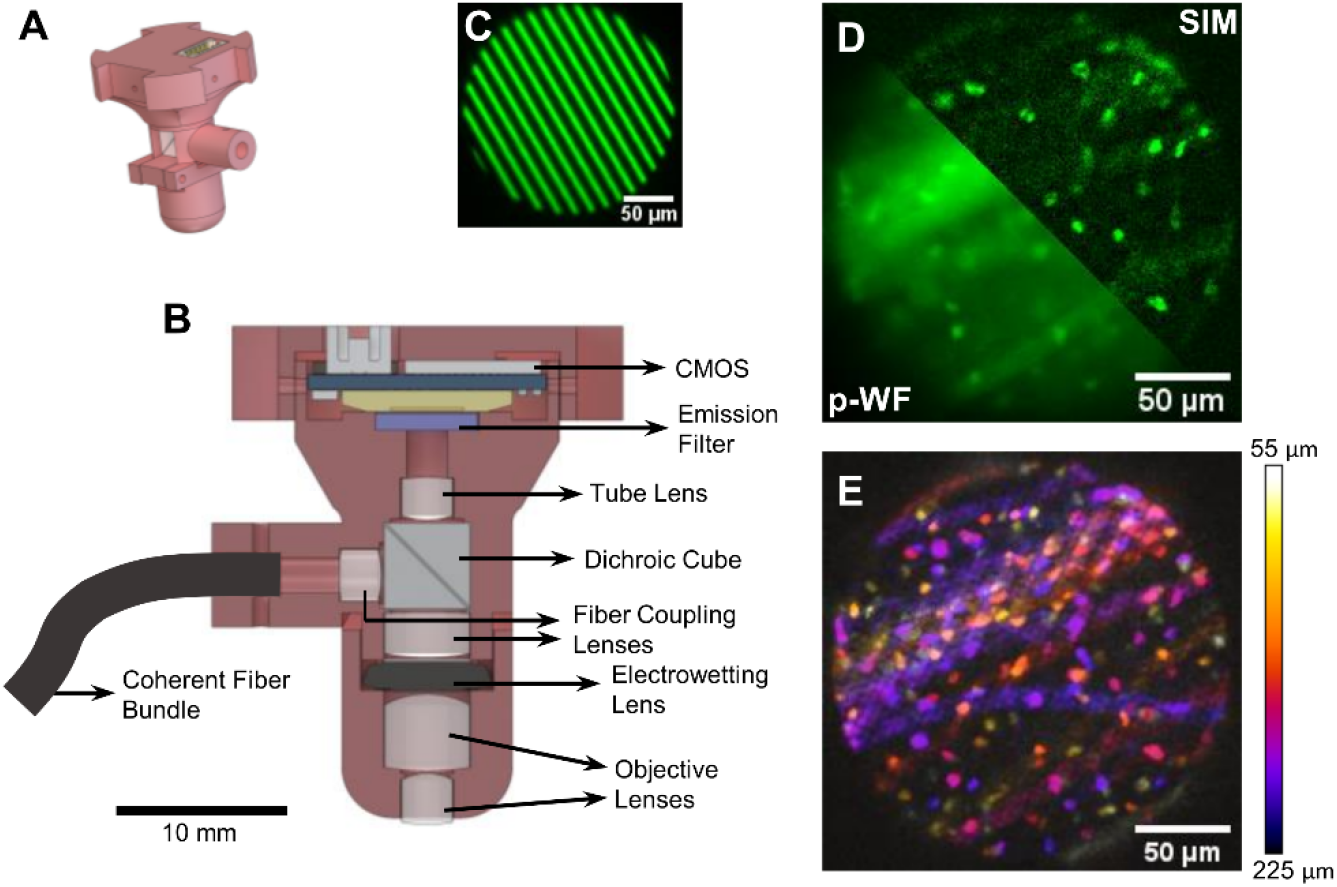
Schematic of the SIMscope3D optical setup. (A) CAD rendering of the SIMscope3D; (B) Cross-sectional view of the SIMscope3D. The design consists of achromatic doublets for fiber coupling and focusing on the sample. The excitation light and fluorescence emission paths are seperated by a dichroic cube. Active axial scanning up to 550 μm into the sample achieved using an electrowetting lens. The fluorescence from the sample is collected onto an onboard CMOS sensor as shown by the stripe pattern shown in (C); (D) SIM reconstructed image compared to the pseuo-widefield (p-WF) reconstruction; (E) Demonstration of volumetric imaging capability of the SIMscope3D with color coded cells from different imaging planes in a fixed tissue sample.

### Imaging sample preparation

For resolution characterization, a sample of fluorescent microspheres (Fluormax G0100) suspended in agarose was prepared in a 35 mm cell culture dish. A coverslip was placed on the exposed surface of the agarose mixture to provide a uniform interface for imaging through. The agarose sample was then mounted under the SIMscope3D for imaging. For fixed tissue imaging, a 350 μm thick coronal slice of fixed, PLP-eGFP mouse brain tissue was mounted on a microscope slide in VECTASHIELD Plus Antifade Mounting Media (H-1900).

For in vivo imaging, cranial windows were implanted as previously described [26]. Briefly, 2 mm^2^ cranial windows were implanted centered on stereotactic coordinates AP +1, ML +1.5 over the motor cortex. A head bar with a 12 mm diameter open window was used to head fix the mouse without interfering with the SIMscope3D. The mice were anesthetized using 4.5% isoflurane and mounted to the head bar apparatus while unconscious prior to imaging. Imaging in identical locations was completed on SIMscope3D and 2-photon systems within 5 hours of surgery. B6.129P-Cx3cr1tm1Litt/J (Jackson lab stock #005582) mice were used for all experiments. All experiments involving animals were conducted in accordance with protocols approved by the Animal Care and Use Committee at the University of Colorado Anschutz Medical Campus.

### SIM Reconstruction

The fluorescent bead, fixed tissue, and live animal images were acquired using three separate grating orientations with three phases per orientation with an incident power of 7.4 μW per phase. The spatial frequency of the grating on the imaging plane was 68.9 mm^-1^ which corresponds to an axial resolution of 17.6 μm with the Stokseth approximation [17]. The exposure time per phase for fluorescent bead and fixed tissue acquisition was 80 ms, and for live animal imaging was 300 ms. The sectioned images for each orientation were obtained using the square law detection method as a SIM reconstruction using Eq. (1) [20]. This step was followed by combining the SIM reconstructed images for each orientation using wavelet fusion [27–29]. Non-sectioned pseudo widefield (p-WF) images were be obtained by adding the images obtained from the three phases and taking the mean intensity of the image over the three orientations.

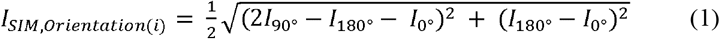

where, *I_0°_, I_90°_* and *I_180°_* are images acquired corresponding to the three phase shifted grid patterns, and *I_SIM,Orientation(i)_* is the SIM reconstruction for the i^th^ orientation.

The p-WF and SIM reconstructed images were processed in ImageJ with brightness and contrast correction, gamma adjust, and a despeckle noise filter. The processing parameters were kept consistent between p-WF and SIM images for every set of images acquired using the SIMscope3D.

## 3. Experimental Results

### Optical Sectioning and Axial Resolution

The optical sectioning characteristics were calculated based on the axial resolution of the SIMscope3D by imaging 1 μm fluorescent beads. The electrowetting lens was set to the center of its actuation range (z = 275 μm, applied voltage of 54 V). A region was chosen with multiple fluorescent microspheres (Figure 2A) in the FOV. Image acquisition for SIM was performed through this region of interest with 61 slices at an axial stepping interval of 1 μm for a total axial depth of 60 μm.

**Figure 2:**
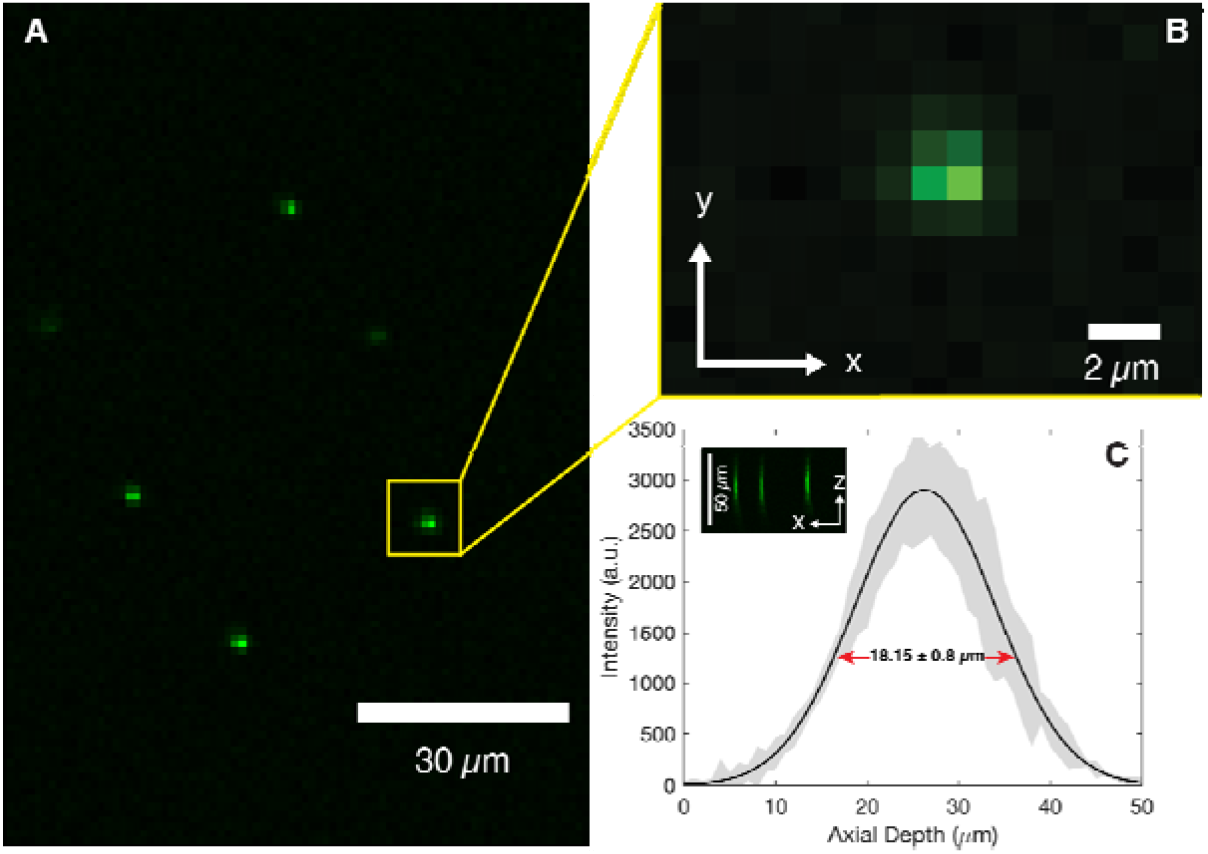
(A) Image of 1 μm diameter fluorescent beads for axial resolution characterization; (B) Close up view of a single bead for characterizing the z-axis profile; (C) The intensity vs axial depth for all fluorescent beads was fit to a Gaussian function, yielding an axial full width half maximum of 18.15 μm. Shaded region corresponds to ±1 standard deviation.

A 25×25 pixel region around each fluorescent bead was cropped through the depth of the image stack (Figure 2B). The intensity profile for each fluorescent bead along the x-axis of each image in the stack was captured using an ImageJ macro. The maximum intensity across each slice was extracted for each fluorescent bead. The maximum intensity vs. axial depth was averaged across all fluorescent beads and fit with a single-term Gaussian function. The axial resolution of the SIMscope3D as indicated by the full width at half maximum (FWHM) of this fit is 18.15 +/- 0.8 μm (Figure 2C).

### Fixed tissue imaging

To demonstrate the viability of the SIMscope3D for biological tissue imaging, we imaged regions in the striatum in a fixed brain tissue labeled with PLP-eGFP. Figure 3 shows a comparison between p-WF and SIM reconstructed images. The effect of optical sectioning and rejection of out of focus fluorescence and scattering background is evident in Figure 3C comparing line cut intensity plots between p-WF and SIM images. Attributed to the higher contrast obtained via optical sectioning, the SIM images can distinguish oligodendrocytes that are buried in the background from nearby bundles of myelinated fibers.

**Figure 3:**
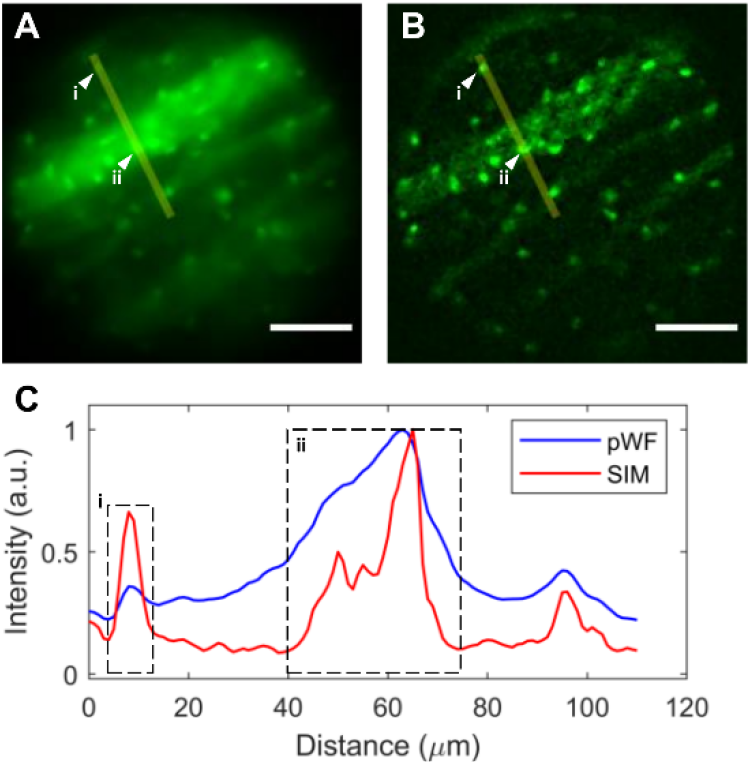
Fixed PLP-eGFP mouse brain tissue imaged with the SIMscope3D. Comparison between the pseudo widefield (A) and SIM reconstructed (B) image is shown; (C) The intensity profile of the line cut indicated in (A) and (B) demonstrating rejection of out of focus background leading to increased contrast and higher visibility of oligodendrocytes in the focus plane. The ability to resolve oligodendrocytes from nearby bundles of myelinated fibers (i and ii) is especially evident. Scale bar: 50 μm.

The SIMscope3D is designed to perform non-mechanical depth scanning up to 550 μm using the integrated electrowetting lens. We performed volumetric imaging with axial stepping interval of 4.58 μm for a total axial depth of 330 μm, demonstrating its volumetric imaging capability. Figure 4 shows the ability of the SIMscope3D to image and distinguish cell bodies up to a depth of 220 μm. The imaging depth beyond 220 μm is in part limited by the incident power in addition to scattering and loss of modulation contrast from the tissue at higher depths.

**Figure 4:**
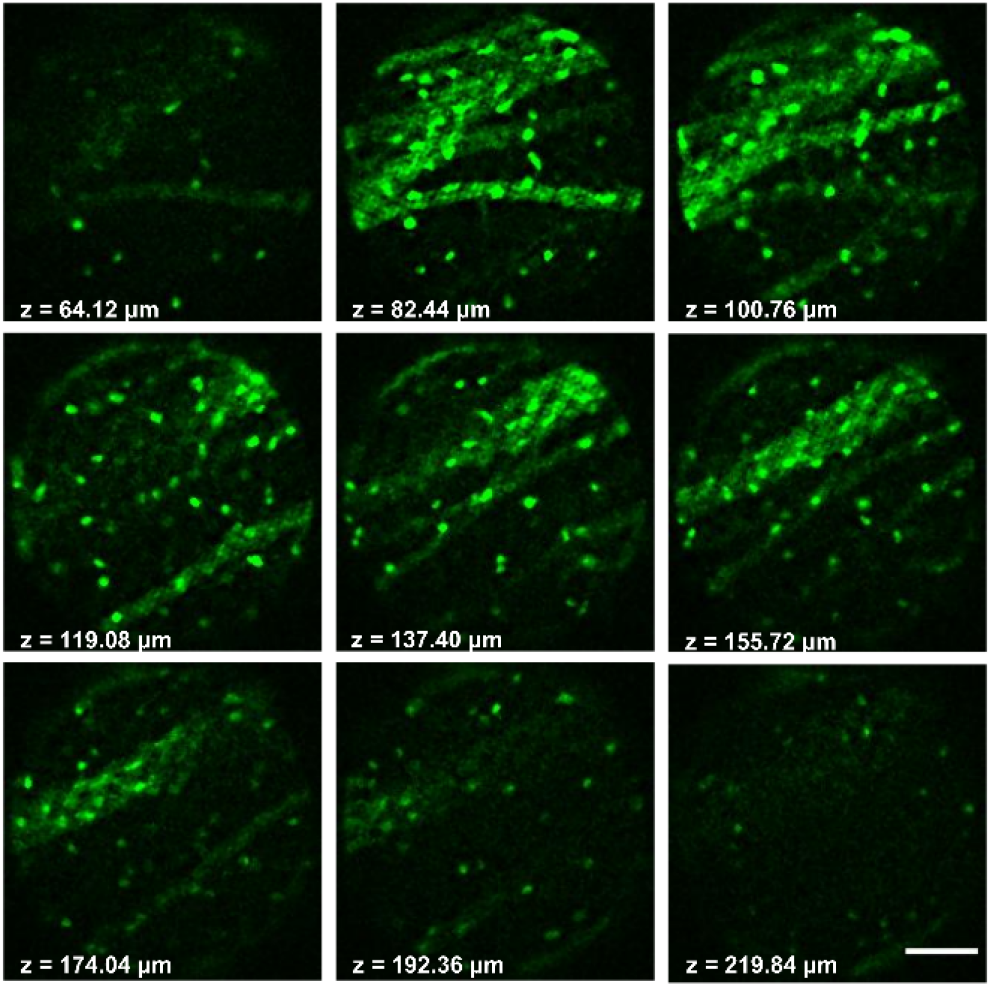
Image stack with increasing imaging depth collected using the SIMscope3D in the striatum region of fixed PLP-eGFP mouse brain coronal slice. With only 7.4 μW/phase incident on the sample, we are able to image and clearly distinguish cell bodies in the sample up to 220 μm deep. Scale bar: 50 μm.

### Microglia processes in awake animals

Microglia, the brain’s tissue resident macrophages, have highly dynamic branched processes that continuously surveil their environment. To image the motion of microglia processes in an awake mouse, the microscope field of view was aligned to a region with visible cells and was positioned next to noticeable vasculature landmarks. The SIMscope3D was focused onto the surface of the cranial window and volumetric images were acquired at five different time points, each volume stack took ~3.5 min with 15 min in between each stack (t = 0, 15, 30, 45, 60 min). The stacks ranged from 0 to 120 um below the bottom of the cranial window with an axial stepping interval of 5 μm. The results from this experiment are presented in Figure 5. The SIM images provide background-free high-resolution imaging of cell bodies even at depth, when compared with p-WF images (Figure 5A). A maximum intensity projection of microglia somas up to a depth of 120 μm are shown in Figure 5B. Figure 5B also demonstrates the ability for the SIMscope3D to optically section cells in densely labeled tissue. The ability to reject out of focus background enables the use of the SIMscope3D to capture a time series of the motion of microglial processes as show in Figure 5C. Additionally, ground truth 2P images were acquired of the same FOV, using the vasculature landmarks as a reference, compared to SIMscope3D images (Figure S4). The 2P images confirmed that the cells being imaged were indeed microglia. In comparison with the p-WF images, the SIMscope3D was able to resolve smaller microglial processes with an improved contrast, where the p-WF image was unable to resolve the feature at all (Figures 5D & 5E). Finally, the SIMscope3D also produces images with greatly improved contrast as compared with the p-WF images (Figure 5F). This proof-of-concept experiment shows that the SIMscope3D can perform long-term awake imaging of microglia, which will allow studies of the dynamic nature of the microglial response in mouse disease models in novel, low-cost ways.

**Figure 5:**
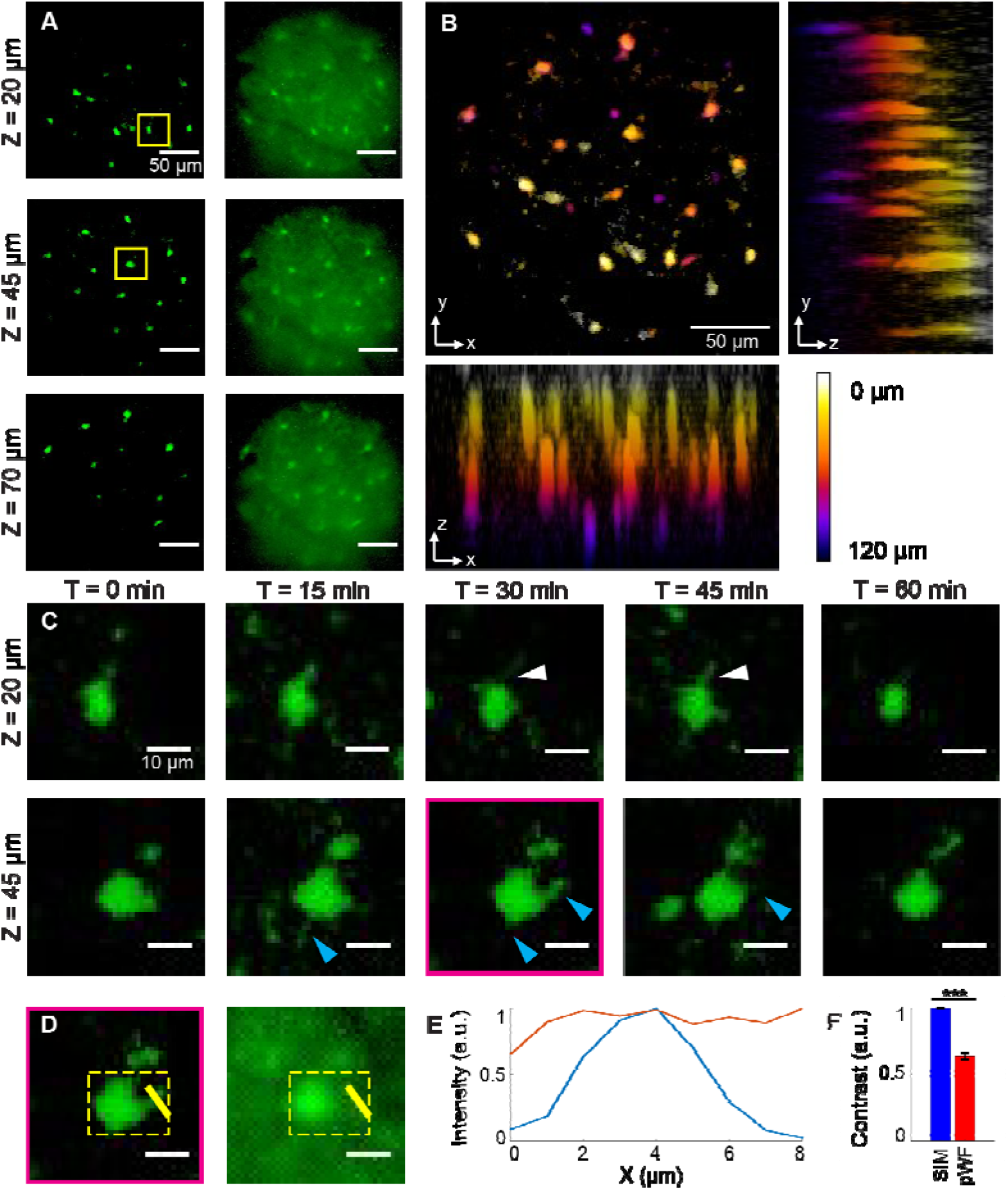
(A) Image stack comparison of microglial cells between SIM reconstructed (left) and pseudo-widefield (right) images, with three different representative z depths (0 μm refers to bottom of cranial window); (B) Color-coded Z-depth max projections for xy, yz and xz planes at T = 0 min; (C) Representative microglial cells across five different time points from ROIs in (A) at depths of 20 μm (top row) and 45 μm (bottom row). Images were averaged across 3 slices (15 μm) for better SNR. White arrows indicate microglial process growth and cyan arrows indicate microglia process retraction; (D) T = 30, Z = 45 ROI from (C) comparing SIM (left) vs. pWF (right); (E) Normalized intensity line profiles for yellow line in (D); (F) Contrast comparison for yellow dotted square ROI in (D) for all five time points (p = .00014, N = 5).

## 4. Discussion

The components chosen in the design of the SIMscope3D, such as achromatic doublets for minimizing chromatic shift, electrowetting tunable lens for depth scanning, and a board level 2.2 μm pixel CMOS sensor, are aimed towards providing high contrast and resolution. This unique benefit makes this miniature microscope different from others that have been designed for imaging neural activity from cells where higher contrast and resolution is not required. In comparison, the SIMscope3D allows us to distinguish feature that are excluded from widefield techniques as shown in Figure 5. This is important for studies of structure changes in supporting cells, such as microglia.

The SIMscope3D also opens new applications in real-time low noise volumetric imaging of neural activities. The neural activity signals obtained by the most popular Inscopix and UCLA miniscopes relies heavily on computation-heavy post hoc imaging processing and noise reduction algorithms. Yet, in densely labeled samples, the neural activity signals obtained by these devices are influenced heavily by out-of-focus fluorescence, thus noise, of neighboring neurons. In applications that require real-time neural signals such as newly emerged calcium-imaging-based brain-machine interface technology [30,31], with slight modifications such as fast frame rate CMOS sensor, the SIMscope3D is capable of providing high quality real-time neural signal while maintaining other advantages of miniscopes.

The main advantage offered by the SIMscope3D is that it can optically section densely labeled scattering tissue (Figure 5B) which improves image contrast (Figures 5E and 5F). This provides the ability to identify prominent cellular features with not only lateral but also axial spatial information, giving researchers the ability to study the 3D structure of neural and non-neuronal populations in real-time. In contrast, conventional miniature widefield microscopes do not offer optical sectioning and have a large depth of field, preventing extrapolation of axial information. An alternative solution for high resolution imaging in freely moving animals is the two-photon fiber coupled miniature microscope systems (2P-FCM). But these systems require bulky, expensive lasers and a more complex optical setup, making the SIMscope3D an easier and more cost-effective system to implement. Thus, the SIMscope3D offers a novel, easy-to-implement way for researchers to perform high-contrast 1-photon 3D imaging on neural populations *in vivo*.

Comparing with 2P imaging, the SIMscope3D currently loses some microglial features and processes. However, the 2P images were highly saturated (Supplementary Figure S4B), which highlights the microglial processes with lower signal in comparison with cell bodies. The current SIMscope3D is limited due to the amount of power incident at the imaging plane from our LED source (7.4 μW/phase). With future iterations of the system, the power can be increased to allow better imaging of small features. Additionally, increasing the excitation power can greatly reduce current exposure times, allowing faster frame rate imaging than from 2P-FCM imaging which is limited by laser scanning.

## 5. Conclusion

In this work, we present a design for a 3D structured illumination fiber coupled miniature microscope with an onboard 2.2 μm CMOS sensor and an active non-mechanical axial scanning lens, capable of imaging at depths up to 550 μm in tissue. This is the first demonstration of a miniature microscope for structured light illumination to acquire volumetric high resolution optically sectioned imaging in live tissue.

We experimentally characterized the axial resolution of the SIMscope3D at 18 μm (FWHM). We demonstrated proof of concept volumetric imaging in fixed brain tissue labeled with eGFP at depths up to 220 μm. The higher contrast obtained in SIM reconstruction due to optical sectioning, enabled time series volumetric images of microglia processes in an awake animal, at depths up to 120 μm.

The SIMscope3D opens new applications in structural volumetric imaging in awake animals with high contrast. The benefits of this system include the lower cost and ability to use higher frame rates than 2P miniature microscopes since images are theoretically only limited by the frame rate of the camera. These features create new opportunities to investigate dynamic neural structure and function in behaving animals.

## Funding

National Institutes of Health (R21 EY029458); National Institutes of Health (R01 NS123665).

## Acknowledgments

The authors would like to acknowledge Tarah A. Welton at Department of Bioengineering, University of Colorado Anschutz Medical Campus for help with the fixed coronal brain tissue. The authors would also like to thank Dr. Mo Zohrabi at Department of Electrical, Computer and Energy Engineering, University of Colorado Boulder for technical assistance with Zemax OpticsStudio modeling.

## Disclosures

The authors declare no conflicts of interest.

## Data availability

Data underlying the results available upon reasonable request.

## Supplemental document

See Supplementary Materials for additional information on the optical design, SIMscope3D characterization, fixed tissue imaging, and ground truth 2P images for comparison of awake animal data.

## Notes

### Competing Interest Statement

The authors have declared no competing interest.

