## Supplementary Figures for "Miniature Structured Illumination Microscope for *in vivo* 3D Imaging of Brain Structures with Optical Sectioning"

**Supplementary Materials**

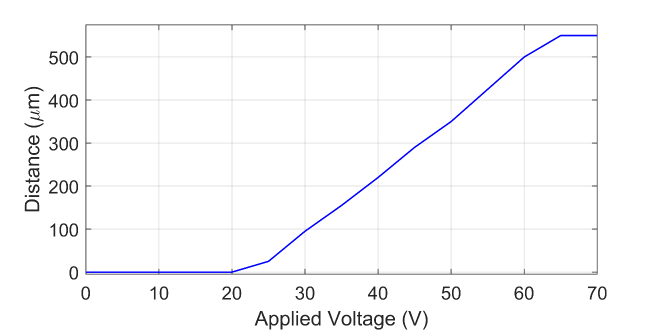

Figure S1: Axial depth calibration of the SIMscope3D as a function of the electrowetting lens applied voltage.

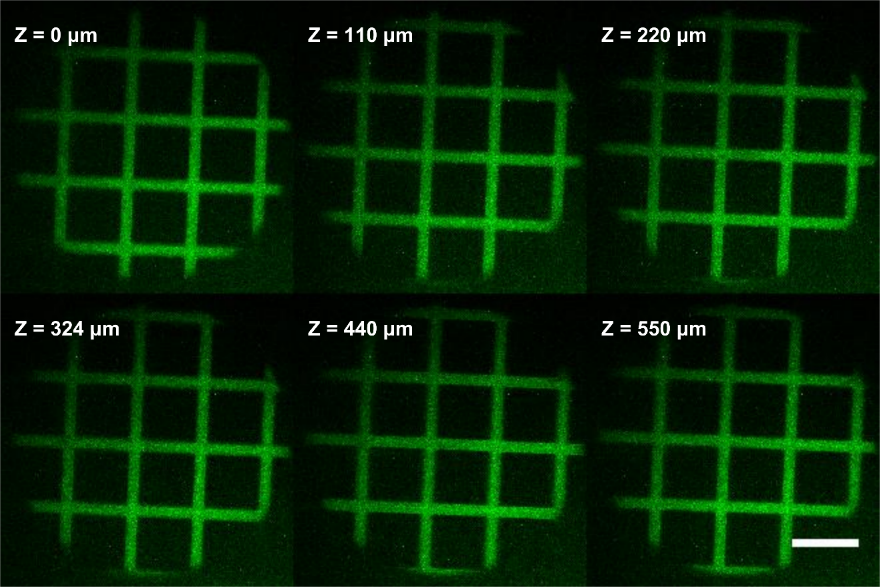

Figure S2: FOV calibration of the SIMscope3D at various axial depths using a 50 μm square grid pattern based on axial calibration from Figure S1. Scale bar is 50 μm.

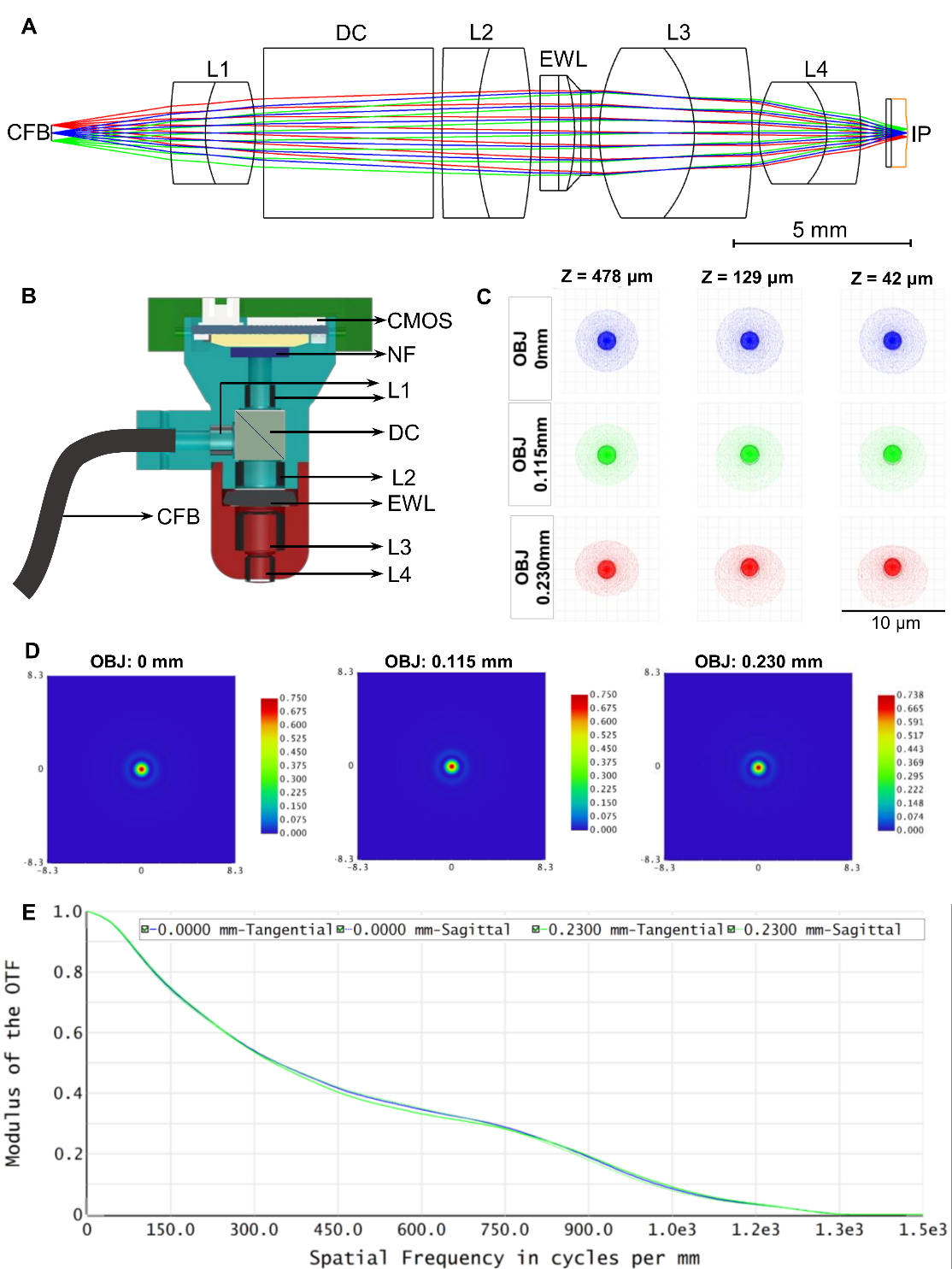

Figure S3: (A) Zemax OpticsStudio design of the excitation path at 470 nm for the SIMscope3D. The dichroic cube (DC) is modeled as a 5 mm path length BK-7 window. The fiber bundle (FB) provides the excitation light for the SIMscope3D. Lenses (L1 to L4) are achromatic doublets, and the electrowetting lens (EWL) provides depth scanning of the imaging plane (IP) of up to 550 μm below the coverslip. The imaging plane has a field curvature of 2.71 mm; (B) Cross-sectional view of the SIMscope3D CAD design with all the components listed. CFB: Coherent Fiber Bundle, Fujikura FIGH-10-500N, L1: Edmund Optics 3mm Dia. x 9mm FL MgF2 coated achromatic doublet lens, L2: Edmund Optics 5mm Dia. x 20mm FL MgF2 coated achromatic doublet lens, L3: Thorlabs 5mm Dia. x 7.5mm FL A coated achromatic doublet, L4: Edmund Optics 3mm Dia. x 4.5mm FL MgF2 coated achromatic doublet lens, DC: Dichroic Cube, Shanghai Optics cemented cube with dichroic coating reflecting 440-480 nm and transmitting 510-550 nm, EWL: Electrowetting Lens, Varioptic Corning A25H, NF: Notch Filter or the Emission Filter, CMOS: CMOS image sensor, Ximea MU9PM-MBRD, IP: Imaging Plane; (C) Full field spot diagram corresponding to object field positions 0mm, 0.115mm and 0.23mm at depths 42μm, 120 μm and 478 μm, here object field position corresponds to the location of the fiber in the CFB; (D) Huygens point spread function at the imaging plane (130μm in the tissue) for various object positions; (E) Huygens modulus of the optical transfer function for various object field positions in the fiber bundle.

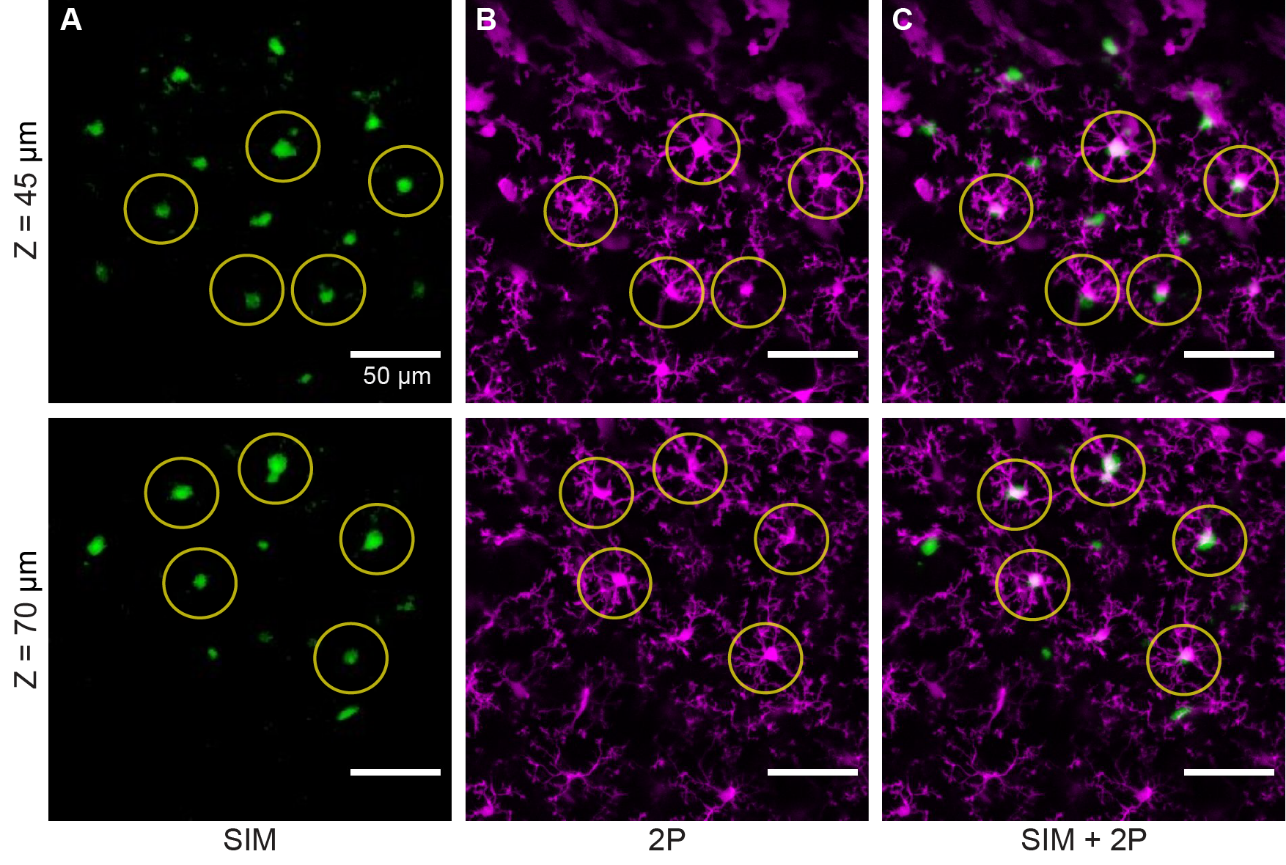

Figure S4: (A) Average Z projection (over 15 μm) of SIM image stack at two depths (top = 45 μm; bottom = 70 μm). The same FOV is used for all panels and ROIs indicate the same cells in each panel at a given depth. (B) Average Z projection of 2P image stack at same depths as A, with same Z depth range. (C) Panels A and B overlaid with areas of direct overlap shown in white. Imperfect overlap may be due to cell migration between imaging. Coverslip was slightly angled for 2P imaging which caused the FOV to be obscured at depths above ~45 μm.
